# Recombinant expression and characterisation of a lipase from the Antarctic zooplankton *Salpa thompsoni*

**DOI:** 10.1101/2022.11.18.517127

**Authors:** Ekta Rayani, Alexander Cotton, Iwan Roberts, John Ward, Will Goodall-Copestake, Brenda Parker

**Affiliations:** Department of Biochemical Engineering, Bernard Katz Building, University College London, Gordon St London, United Kingdom WC1E 0AH; Cytiva Ltd, Stevenage Bioscience Catalyst Gunnels Wood Road, Stevenage, United Kingdom, Hertfordshire, United Kingdom SG1 2FX; Scottish Association for Marine Science, Oban, Argyll, Scotland PA37 1QA

**Keywords:** Lipase, salp, bioprospecting, biocatalysis, psychrophilic enzymes

## Abstract

Cold marine environments are abundant on earth and represent a rich resource for low temperature enzymes. Here we apply *in silico* bioprospecting methods followed by *in vitro* expression and biochemical analyses to characterise a novel low temperature lipase from the Antarctic tunicate *Salpa thompsoni*. A 586 amino acid pancreatic lipase-like gene was identified from *S. thompsoni* transcriptomic data, expressed as a hexahistadine fusion protein in *Escherichia coli* at 10°C and purified by affinity chromatography. Hydrolysis of the synthetic substrate ρ-nitrophenyl butyrate (PNPB) showed that this recombinant protein has optimal activity at 20 °C and pH 7, and a specific activity of 3.16 U/mg under this condition. Over 60% of enzyme activity was maintained between 15 to 25 °C, with a sharp decrease outside this range. These results are indicative of cold active psychrophilic enzyme activity. A meta-analysis of lipase activities towards PNPB showed that the novel *S. thompsoni* lipase displays a higher activity at lower temperatures relative to previously characterised enzymes. The work demonstrates a methodology for conversion of transcriptomic to *in vitro* expression data for the discovery of new cold-active biocatalysts from marine organisms.

## Introduction

Low temperature environments, while seemingly inhospitable, contain a variety of life forms that present a rich treasure trove for bioprospecting (Wagner Fernandes Duarte et al. 2018). The world’s oceans that encompass the largest biome on Earth (Keith et al. 2020) and most cold environments are marine. These include not only polar surface waters but also waters found at depth below the thermocline. Despite the broad geographic coverage of low temperature marine environments, they remain underexplored from a bioprospecting perspective because they are typically remote and complex to access.

Psychrophilic biocatalysts are cold adapted enzymes that may be found in organisms inhabiting environments where the temperature for growth is 15°C or lower (Moyer and Morita 2007). By definition, these enzymes are most active below 30°C (D’Amico et al. 2003; Feller and Gerday 2003). Psychrophilic enzymes possess several features that are thought to ameliorate the exponential slowing of chemical reactions at decreasing temperatures to maintain crucial functions. Glycine clusters around the active site provide thermostability and structural flexibility to encourage higher activity (Veno et al. 2019). Increased hydrophobic surface residues and increased hydrophilic groups within protein cavities contribute to conformational stability and enhance ionic reaction catalysis (Metpally and Reddy 2009). Amino acid composition also influences the overall conformational stability, for instance a high lysine-to arginine ratio was previously shown to be important in cold adaptation (Siddiqui et al. 2006).

Enzymes from the triglyceraldehyde hydrolase family, known as lipases, E.C 3.1.1.3, are an industrially important class of biocatalysts. Their primary function in native conditions is to hydrolyse carboxylic ester bonds in hydrophobic compounds. In aqueous conditions lipases catalyse the hydrolysis of carboxylate ester bonds into free fatty acids (FFAs) and organic alcohols. Lipases are also able to catalyse reactions in organic solvents including esterification and transesterification (Chandra et al. 2020). A range of recombinant lipases have been produced from bacterial (Salwoom et al. 2019; Schumann and Ferreira 2004; Guozeng Wang et al. 2016), fungal (Lin et al. 2017; S.-J. Tang et al. 2000) and mammalian sources (Kawaguchi et al. 2018; Valdez-Cruz et al. 2017). These industrially relevant classes of enzymes include *Candida antarctica* Lipase B (Cal-B) derived from a low temperature Antarctic yeast. This enzyme is valuable due to its high enantioselectivity, ability to recognise a wide range of substrates, thermal stability, and capacity to catalyse reactions in polar solvents which facilitates the application of Cal-B in organic synthesis (Stauch, Fisher, and Cianci 2015).

Lipases that have been optimised to function at lower temperatures have potential applications in the fine chemical and personal care sectors. They have been applied for the synthesis of high value chiral intermediates for pharmaceutical applications (Patel, Banerjee, and Szarka 1997) and for the production of biologically active compounds such as xanthenes used in the synthesis of insecticidal molecules (Jiang et al. 2014). Furthermore, low temperature lipases afford the opportunity for enzyme inactivation at higher temperatures to enable precise control over thermally sensitive reactions (De Santi et al. 2014). For bulk processes, the lower energy requirement of cold temperature biocatalysts can represent a benefit in terms of sustainability (Cavicchioli et al. 2011). This range of applications has motivated explorations for novel lipases from multiple environmental sources. However, while lipases from polar bacteria and yeasts have been explored for low temperature activity, lipases originating from more complex eukaryotic organisms inhabiting cold environments remain under-investigated as potential sources for new biocatalysts.

*Salpa thompsoni* is a marine animal found in the Southern Ocean surrounding Antarctica (Figure 1). Commonly referred to as a salp, it is a zooplankton from the tunicate group, which is an early diverging lineage of chordates (Ambrosino et al. 2019). In contrast to the vast majority of salp species that occur in tropical to temperate waters, *S. thompsoni* appears to have evolved as a low temperature adapted species (Goodall-Copestake 2018). It is capable of immense population blooms at temperatures from 3°C to 5°C degrees (Henschke and Pakhomov 2019). As such, this species might be expected to have evolved an effective low temperature metabolism which potentially involves psychrophilic enzymes of industrial relevance. These could include FFA digesting lipases that assist *S. thompsoni* in processing the wide range of food particles it obtains through filtration (von Harbou et al. 2011).

**Figure 1:**
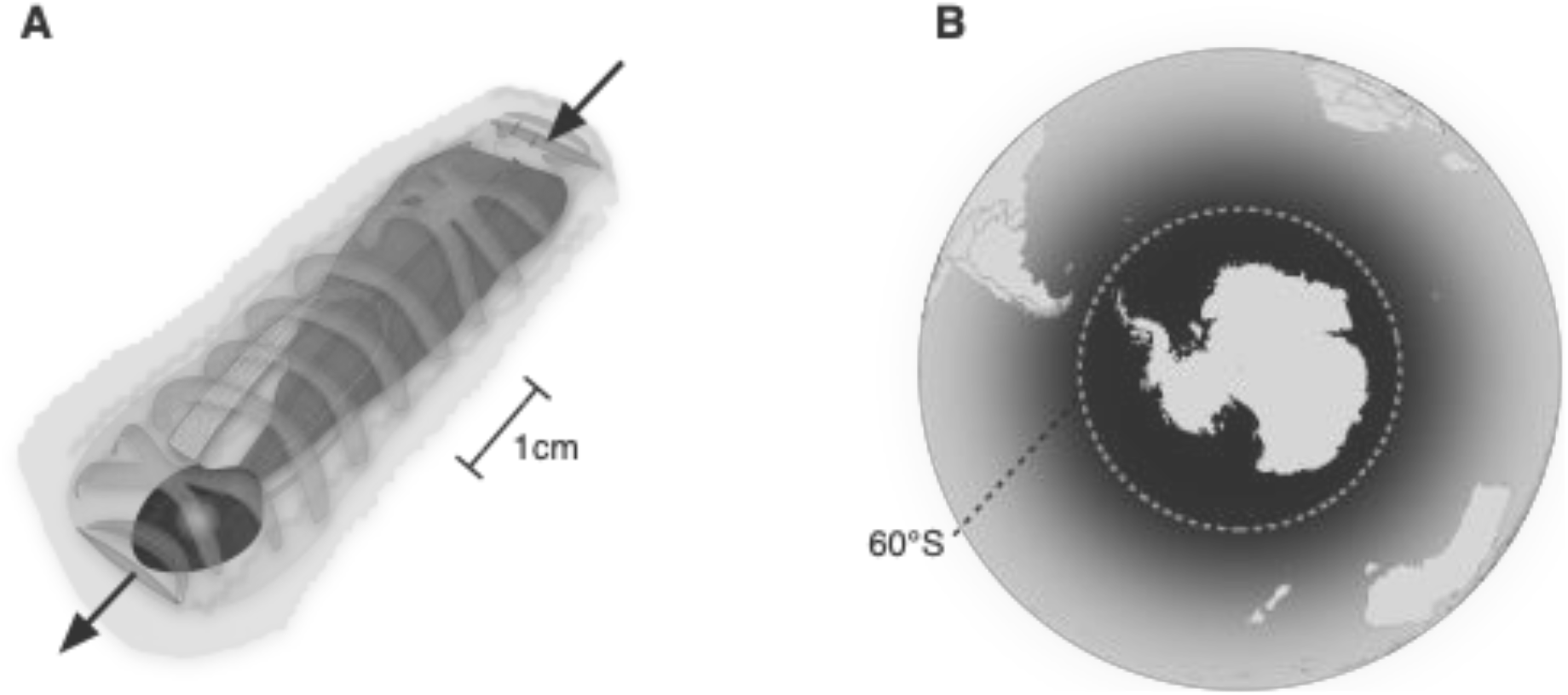
**Schematic of the translucent zooplankton** *Salpa thompsoni* **with gut highlighted in dark grey, filter in mid grey and arrows to indicate water inlet and outlet (A), and generalised** *S. thompsoni* **distribution highlighted in dark grey, where 60 °S marks the northern limit of the Southern Ocean. Figure adapted from** Foxton 1966; Van Soest 1974.

Here we used a bioinformatic bioprospecting workflow to identify a putative lipase gene from *S. thompsoni* as a novel, potential low temperature lipase. This was subsequently expressed in a bacterial host and purified to allow for functional characterisation using a common substrate. Optimising heterologous expression in bacterial host systems, such as *Escherichia coli* facilitates more robust manufacture of biocatalysts (Rosano and Ceccarelli 2014). The results obtained were compared against previously published activities on the same substrate from other animals, plants, fungi and bacteria in order to contextualise cold temperature activity amongst lipases from all kingdoms of life.

## Materials and Methods

### Identification

A putative *S. thompsoni* lipase was identified using the annotated genome of the tunicate and model organism *Ciona intestinalis* (Godeaux, 1981). NCBI GenBank was interrogated using the search terms “Ciona intestinalis”[Organism] AND lipase [All Fields] (Agarwala et al. 2018; Altschul 1997) to recover a ‘pancreatic lipase-related protein’ from *C. intestinalis*; GenBank accession **XP_002130573.1**. As pancreatic lipases are one of the main digestive enzymes used to break down dietary fat molecules in animals (Zhu et al. 2021), this accession was used to search for a novel, orthologous lipase gene from *S. thompsoni*.

The un-annotated *S. thompsoni* transcriptome assembly (GenBank **GFCC00000000.1**; Batta-Lona *et al*., 2016) was queried using the accession **XP_002130573.1** with the tBLASTn tool as implemented in the software package GeneiousPrime version 2021.1 (Kearse *et al*., 2012). This identified *S. thompsoni* assembly sequence **GFCC01067991** which contains a 1761bp open reading frame and putative orthologue of **XP_002130573.1**. To assess if this enzyme was expressed in the digestive system, short read archive data generated solely from *S. thompsoni* gut RNA (samples AC_e20 and AC_e242, accession numbers **SRX13276964** and **SRX13276963**, respectively) were screened for matching sequences using the Map to Reference method in Geneious under the LSF setting (word length 24, index word length 14, maximum mismatches per read 10). Sequence characterisation of the 1761bp lipase sequence was carried out using the Expasy Protein Parameter tool to identify physical and chemical parameters including the molecular weight and amino acid composition (Artimo P 2020; Gasteiger et al. 2005). InterProScan analysis was also used, to identify predictive domains using a genome-scale protein function classification (Jones et al. 2014).

### Expression of lipase

The 1761bp *S. thompsoni* lipase sequence was synthesised and codon optimised by GenScript (Leiden, Netherlands). Codon optimisation increased the GC content from 44.4% to 55%. The codon-optimized lipase was cloned within the expression vector pET-28a (+) with an N-terminal hexahistadine tag to aid purification. The plasmid was transformed into competent *E. coli* ArcticExpress (DE3) cells (Agilent, Santa Clara, CA, USA) through heat shock methodologies following manufacturer’s guidelines. Two biological chaperonins from cold active species were co-transformed for increased protein solubility, Cpn 10 and Cpn 60, with high similarity to the *E*.*coli* chaperones GroEL and GroES (Belval et al. 2015; Ferrer et al. 2003). *Escherichia coli* harbouring the salp lipase was inoculated overnight in terrific broth (TB) medium (10ml) supplemented with kanamycin and gentamycin and grown at 37°C and 125 rpm. Subsequently, 500 mL TB medium was inoculated with 2 mL of the preculture and incubated at 30°C until an OD600 of 4 was reached. Gene expression was induced on ice by addition of isopropyl-β-D-thiogalactopyranoside (IPTG) to a final concentration of 1mM. The cultures were incubated at 10°C at 180 rpm for 24 h. Cultures were harvested after 24 hours by centrifugation (10,000g, 10 minutes at 4 °C). Cell pellets were collected and stored at -20°C for further analysis (Akbari et al. 2010).

Cultures were analysed for expression levels using Bolt™ Sample buffers and Reducing Agents as per the manufacturer’s guidelines, with 12% gels stained using Coomassie brilliant blue for SDS analysis. Gels were visualised using Amersham™ Imager 600 (GE healthcare, USA). Western blotting (Burnette 1981) was carried out using the Bio-Rad Trans-Blot® TurboTM Transfer System. Subsequent methodologies were adapted from previous work (Colant et al. 2021). The membrane was incubated with the primary antibody: Anti-His mouse monoclonal antibody (ab18184, Abcam) diluted 1:1000 in TBST-M for two hours. The membrane was then washed with TBST before being incubated with the secondary antibody: Goat anti-mouse HRP conjugated antibody (HAF007, Biotechne), diluted 1:1000 in TBST-M. Imaging took place using The Pierce ECL Western Blotting Substrate (Thermo Fisher Scientific) for 1-2 minutes in darkness then exposed and imaged using an Amersham™ Imager 600 (GE Healthcare).

### Purification

Purification was carried out as described by (Spriestersbach et al. 2015). Ni-NTA Superflow cartridges (Qiagen, Sussex, UK) were equilibrated with 10 mL of binding buffer (50 mM NaH_2_PO_4_, 300 mM NaCl and 10 mM imidazole). The clarified lysate was loaded into the cartridge and subsequently washed with 20 mL of wash buffer (50 mM NaH_2_PO_4_, 300 mM NaCl and 25 mM imidazole). The enzyme was eluted with 5mL of elution buffer (50 mM NaH_2_PO_4_, 300 mM NaCl and 150 mM imidazole). Ammonium sulfate at 70%, was used to precipitate and store the protein at 4 °C.

### Characterisation

A lipase activity assay was performed using ρ-nitrophenyl substrates with modifications (U. K. Winkler and Stuckmann 1979). In brief, the reaction mixture consisted of 125 µl of 55 mM phosphate buffer adjusted to pH 7 and 10 µl of the substrate 1 mM ρ-nitrophenyl butyrate (PNPB) (Sigma N9876) homogenised in 2-propanol. The mixture was pre-incubated at 20°C for 10 minutes and then 15 µl of the 0.5mg/ml enzyme solution was subsequently added. After 15 minutes of incubation at 20°C, the reaction was terminated using an equal volume of 0.25 M sodium carbonate pH 10.5 (Ozcan et al. 2009) and ethanol. The absorbance of the liberated 4-Nitrophenol was measured at 410 nm using multi-well plate reader (Clariostar, BMG Labtech). One unit (U) of lipase activity was defined as the amount of enzyme that released 1 µmol of 4-Nitrophenol per minute. Lipase concentration was measured by the method of Bradford with bovine serum albumin as a standard (Bradford 1976).The temperature profile of the salp lipase was measured at 5 to 60 °C for 15 minutes. A pH stability test was performed by pre-incubating the lipase for 10 minutes at 4 °C in the buffers and assessing activity at the optimum temperature. The effect of pH on lipase activity was assayed at pH 6–9 by measuring the amount of 4-Nitrophenol liberated using PNPB (C4) as a substrate. The buffer systems used to assess pH effects were: 55mM sodium phosphate buffer (pH 6–8), 55mM Tris–HCl buffer (pH 9).

## Results and Discussion

A putative 1761bp salp lipase, subsequently referred to as PL002, was identified within the salp transcriptome assembly sequence GFCC01067991 using bioinformatic methods. This sequence had a 46% amino acid identity to the pancreatic lipase accession XP_002130573.1 from *C. intestinalis*, and a 36% similarity to that of the Human Pancreatic Lipase (Waterhouse et al. 2018) inferred through the Swiss Model Homology tool, thus suggesting that PL002 is a pancreatic-like lipase (Figure 2). Identical DNA sequence transcripts were found within cDNA reads isolated from the *S. thompsoni* gut sample AC_e20, and 99.7% identical transcript sequences were recovered from *S. thompsoni* sample AC_e242. This corroborated the identification of this lipase as of salp origin and demonstrated that it is expressed in the gut consistent with the hypothesised digestive function.

**Figure 2:**
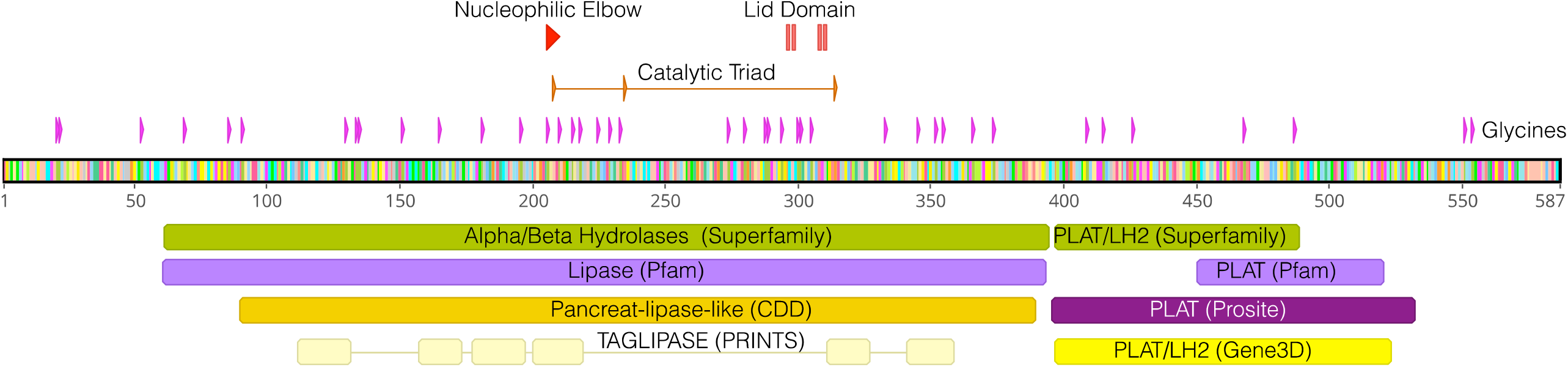
Annotated PL002 amino acid gene sequence of the putative pancreatic salp lipase with InterProScan identified motifs. PLAT (Polycystin-1, Lipoxygenase, Alpha-Toxin) domains, Lipase domains, LH2 (Lipoxygenase homology) identified using InterProScan analysis. Further structural motifs and molecular functions have been highlighted.

Bioinformatic characterisation of sequences was used to confirm several structural features found within lipases. Alpha-beta hydrolases locate the nucleophilic serine across the two other residues of the catalytic triad (Denesyuk et al. 2020). The catalytic triad consists of Ser-208, from the conserved pentapeptide, Asp-235 and His-314 (Figure 2), which is critical for lipase activity (Brumlik and Buckley 1996; F. K. Winkler, D’Arcy, and Hunziker 1990) and common to all enzymes from the serine hydrolases family (Lazniewski et al. 2011). It was found in PL002 alongside the conserved pentapeptide, Gly-206 Phe-207 Ser-208 Leu-209 Gly-210, corresponding to a nucleophilic elbow motif housing a nucleophile serine residue required for catalytic activity (Figure 2). A SignalP-TM was also identified in the N-terminus which may suggest that this lipase is in the secretory pathway but may not necessarily be a secretory enzyme (Nielsen 2017).

Another highly conserved feature of lipases present within the PL002 sequence that occurs in the catalytic domain was Cys-295, His-296, Val-311 and Cys-312. Variations in this lid domain can be used to explain substrate affinity (Ollis et al. 1992; Saavedra et al. 2018) and temperature dependence studies have shown that the lid structure in cold active lipases is smaller than in other lipases active at higher temperatures (Khan et al. 2017). A similar lid structure has also been identified in other aquatic invertebrate neutral lipases from the microcrustacean *Daphnia pulux* (Colbourne et al. 2011). It has been suggested that the cysteine residues contribute towards a disulfide bridge, this has been observed in mammalian pancreatic lipases, particularly that of the secondary fold, the β9 loop (Lowe 2002).

The Expasy Protein Parameter tool (Duvaud et al. 2021) showed that 50% of all glycine amino acids were clustered around the nucleophilic elbow and the active site/lid (Figure 2). In this region glycine clusters potentially encourage greater local mobility of the active site and also support the stability of the enzyme (Mavromatis et al. 2002; Siddiqui et al. 2006).

InterProScan analysis predicted domains within the salp lipase (Lu et al. 2020) based on conserved molecular functions (Figure 2). They included the PLAT (Polycystin-1, Lipoxygenase, Alpha-Toxin) and LH2 (Lipoxygenase homology) domains that are found within membrane or lipid associated proteins (Bateman and Sandford 1999). These domains were identified from Superfamily (Gough et al. 2001; Wilson et al. 2009) and Pfam (Mistry et al. 2021) databases. Lipase domains were also identified through PRINTS (Attwood et al. 2003), Pfam (Mistry et al. 2021) and CDD (Lu et al. 2020) protein prediction software databases.

Purification of PL002 was aided by recombinant production in the Arctic Express cell line (Figure 3A and C). The visualization of purified protein was complicated by the similarity of the size of PL002 at 65kDa and the ArcticExpress (DE3) cell line chaperone, Cpn 2 at 57 kDa. Due to this size similarity, a His-tag Western blot analysis was employed to confirm the presence of the protein (Figure 3B). Pure elution of PL002 was ensured by increasing imidazole concentration of the binding buffer by 10mM, as demonstrated by previous separation studies (Hartinger et al. 2010). The overall recovery of soluble fractions purified in one step affinity chromatography was 60%.

**Figure 3:**
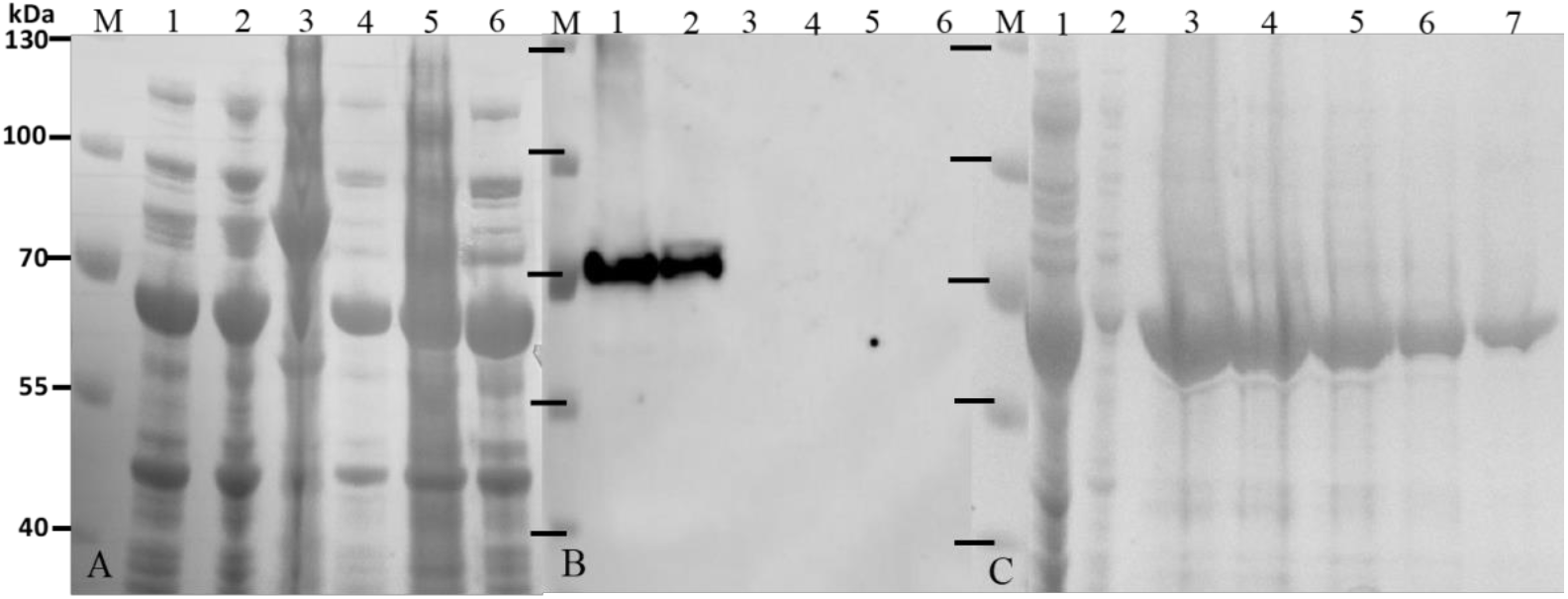
Confirmation of recombinant expression of PL002 using SDS-PAGE (A), Western blot (B), and Purification of PL002 (C) A Coomassie Blue stained SDS PAGE (A) and its corresponding Western Blot developed using chemiluminescence with the Anti-His mouse monoclonal antibody (ab18184, Abcam) For images A and B: Lanes 1 and 2: PL002 clarified lysate and insoluble fraction Lanes 3 and 4: control pET-28a (+) empty vector clarified lysate and insoluble fraction Lanes 5 and 6: control untransformed ArcticExpress (DE3) clarified lysate and insoluble fraction For image C: Lane 1 and 2: Flowthrough and wash fraction (50 mM) of PL002. Lane 3-7: imidazole elusion steps (150 mM)

Under the conditions evaluated, enzyme characterisation using the PNPB substrate showed a peak in activity for PL002 at 20°C and pH7 (Figure 4A and B). It was observed that 60% of relative activity was retained between 15 and 25 °C, however PL002 loses close to 90% of its activity at temperatures greater than 50 °C which implies that this lipase meets the definition of cold active (Gatti-Lafranconi et al. 2010). Lipases isolated from other eukaryotic sources have been identified to possess a similar pH optimum within the range of 6.2 and 7 (van Kuiken and Behnke 1994). This pH condition is crucial to maintaining the form of the lipolytic lid in lipases, which is composed of charged residues required for substrate binding.

**Figure 4:**
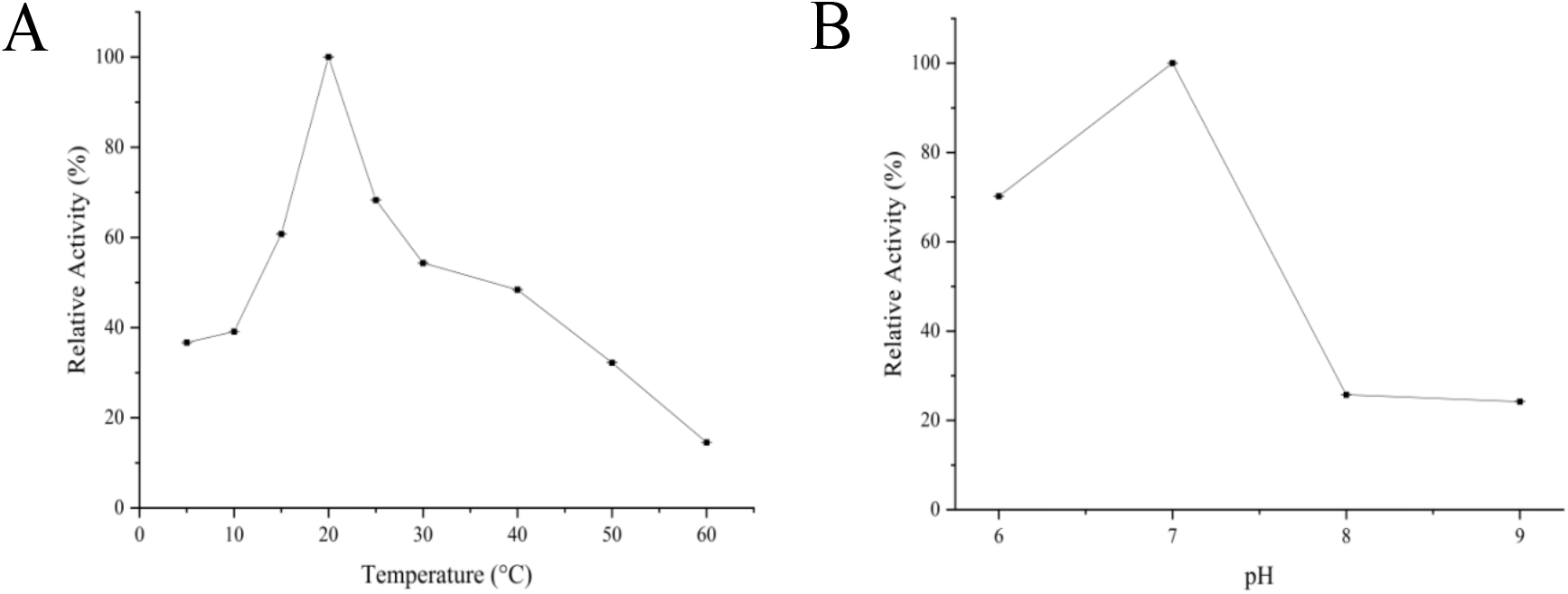
The effect of temperature (A) and pH (B) on PL002 lipase activity on the PNPB substrate. Each data point represents the mean ± standard deviation (SD) of five replicate assays. Activity profiles were expressed in relation to the maximum value (=100%).

The highest activity using PNPB synthetic substrate measured for PL002 was 3.16 +/-0.14 U/mg under conditions of 1mM of PNPB, 20 °C and pH 7. To understand this in relation to other enzymes, a meta-analysis was conducted to compare reported lipase activity against the PNPB substrate in a range of psychrophilic, mesophilic, and thermophilic enzymes (Figure 5). In comparison to other eukaryotic lipases, PL002 exhibited a higher activity on this substrate and the lowest optimum temperature (Supplementary Table 1.3). This is consistent with the evolutionary adaptation of *S. thompsoni* to life in a low temperature marine environment and highlights the potential of this organism as a source of novel psychrophilic biocatalysts.

**Figure 5:**
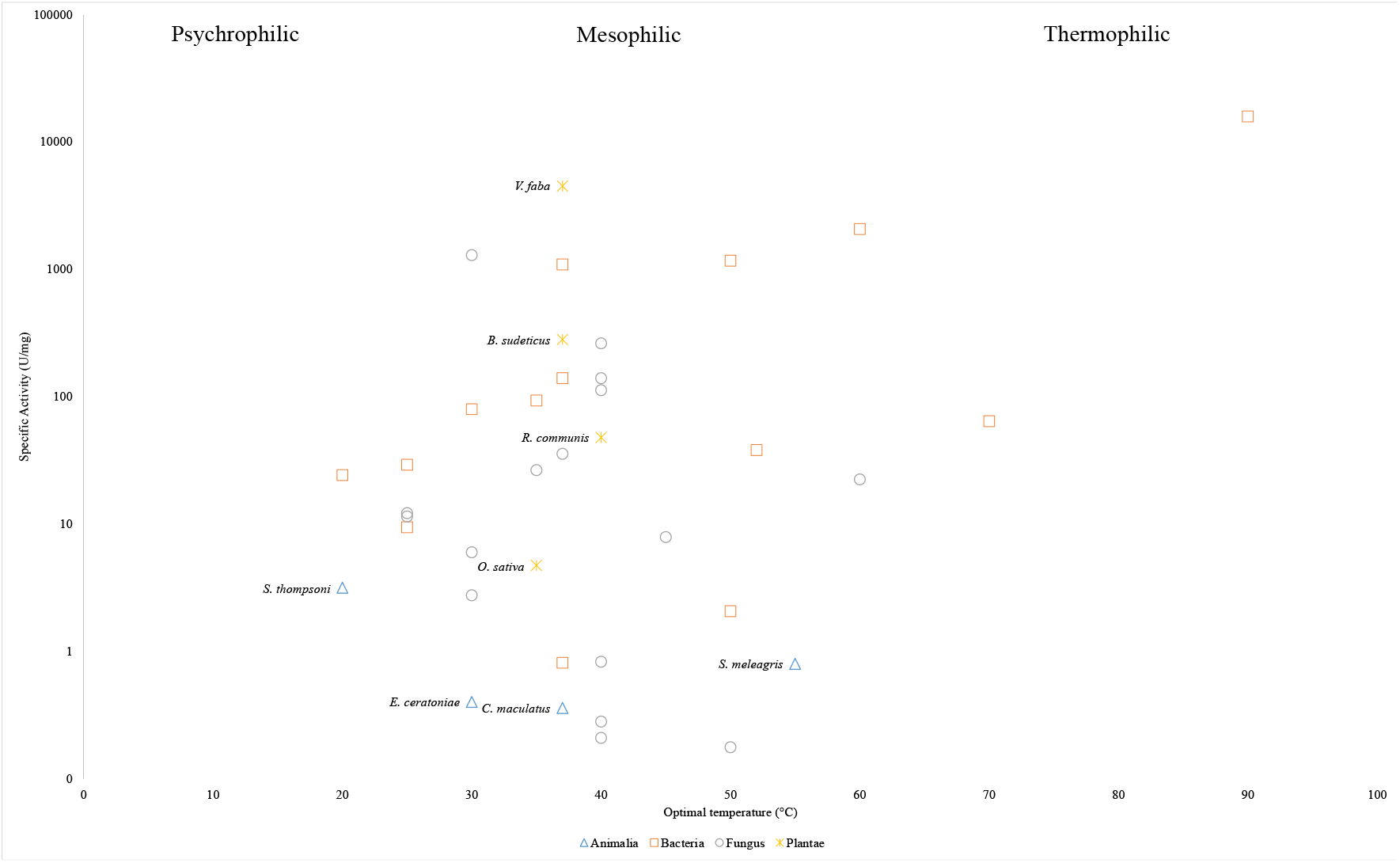
Specific activity (U/mg) of PNPB active lipases from Bacteria, Animalia, Fungus and Plantae. Temperature domains adapted from (D’Amico et al. 2003). Animalia and plantae lipases are labelled in the figure. The origin of each lipase and its class are detailed in Supplementary information Table 1.3

Furthermore, this result provides an additional putative example of a genomic change in *S. thompsoni* in response to low temperatures, adding to reports of increases in nuclear ribosomal DNA content in this species that may represent an adaptation compensating for slower metabolism at low temperatures (Goodall-Copestake 2018; Jue et al. 2016). Subsequent temperature studies with PL002 could include the equivalent lipase from a salp species found in a more tropical climate. This will encourage further refinements of *in silico* bioprospecting whilst functionally characterising eukaryotic cold candidates.

Marine invertebrate genomes have been mined to uncover multiple lipase genes that display different specificities towards different substrate sizes. Studies on the lipid metabolism of the whiteleg shrimp *Penaeus vannamei* (Rivera-Pérez and García-Carreño 2011) found optimal enzyme activity when substrates of a similar nature to their feed source were used, compared to the synthetic substrate. Hydrolysis of neutral lipids is an important aspect of organismal development, and many organisms possess lipase genes defined as functionally hydrolysing triacylglycerol substrates (Rivera-Perez 2015). A complex suite of such enzymes probably works in tandem for digestive purposes. Indeed, it is conceivable that *S. thompsoni* may have a more specialised lipase profile, such as that found in the gastric pouches of the jellyfish (*Stomolophus sp. 2*) where, in comparison to lipases from other invertebrates, lipolytic enzymes favour shorter and medium length triacylglycerides (Martínez-Pérez et al. 2020). The size of carboxyl ester based substrates hydrolysed has been used to categorise enzymes candidates as either a lipase or esterase. A more complete classification of lipolytic esterases should also investigate the ability to hydrolyse long chain fatty acids (Ali, Verger, and Abousalham 2012). Generally, it has been identified that true lipases hydrolyse longer chain fatty acid-based substrates while esterases act on shorter chain substrates, such as PNPB (De Santi et al. 2014). Multiple lipase candidates may exist in eukaryotic invertebrates therefore the activity profile generated from *in vitro* characterisation can be used to describe enzyme function and activity against a wide range of substrates (Volokita et al. 2011). An alternative bioprospecting approach may be to use proteomic and metabolomic trials to understand the lipid preferences of enzymes *in vivo* to uncover industrially promising candidates.

While the activity for PL002 using synthetic substrate, PNPB, was within detectable limits, the impact of biological modifications such as the addition of affinity tags on enzymes, including hexahistadine tags, have been shown to diminish PNP ester hydrolysis activity (de Almeida et al. 2018). It has been previously demonstrated for a *Staphylococcus aureus* lipase that an N terminal His tag may alter the specific activity, the chain length selectivity and the thermostability (Horchani et al. 2009). In another case, it was reported that there was a 33% increase in specific activity as well as preference towards medium length chain fatty acid pNP-laurate (C12) of an untagged recombinantly produced lipase from *Geobacillus kaustophilus* (Özdemir, Tülek, and Erdoğan 2021). There has also been evidence which indicates that this hexahistadine tag can alter the quaternary structure of proteins (Carson et al. 2007) impacting their specific activities. As such, it is plausible that such a modification may have altered the activity of the recombinantly produced affinity tagged protein PL002. Subsequent analyses could explore this further through a comparison against histidine cleaved proteins generated at both the expression and characterisation stages of *in vitro* enzyme production.

To expand the toolkit for low temperature biocatalysis, the pipeline demonstrated here for the identification of cold active lipase candidates using *in silico* methods can be followed. Here we have employed this workflow to identify a pancreatic-like lipase from assembled genomes constructed on sequence homologies of annotated genes. For prokaryotic organisms, culture-independent methods are already a powerful means of overcoming practical limitations in enzyme identification (Sysoev et al. 2021). The characterisation of PL002 from *S*.*thompsoni* is a step towards a wider understanding of how the expanding database of eukaryotic sequences from extreme environments can be a source of promising biocatalysts for temperature-specific industrial applications.

## Summary

This study combined *in silico* bioprospecting with the *in vitro* expression of a gene of interest to characterise a novel source of biocatalytic activity. To our knowledge, this is the first time such an approach has been used to characterise a lipase from a marine tunicate. The lipase was identified from *S. thompsoni* and exhibited psychrophilic characteristics in accordance with the cold Antarctic marine habitat of this organism, retaining over 60% activity between 15 to 25 °C and optimal conditions of pH 7. In future, bioprospecting of the genomic and transcriptomic data available on digestive enzymes from marine invertebrates inhabiting low temperature environments may yield biocatalysts with novel functionality.

## Acknowledgements

ER was funded through an EPSRC studentship (Grant reference number: EP/R512400/1). BP and WGC acknowledge financial support for initial salp gene expression work from the EPSRC Centre for Nature Inspired Engineering (EP/K038656/1)

## Abbreviations

Salps: *Salpa thompsoni*

