## Supplementary Material for "Recombinant expression and characterisation of a lipase from the Antarctic zooplankton *Salpa thompsoni*"

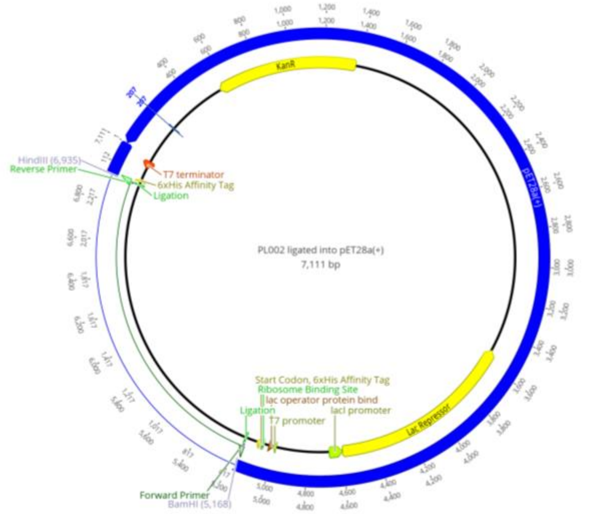


**Figure A: Plasmid map of the Salpa thompsoni Lipase, PL002**

LCAVTVKMTYSSKPSSNNMSGGKSYTWTCSAFLKIILLTLTISPSKAAHPRVGKNETNNK
KEVCYKTYGCFKNDPPFNPKMPLPLGEKQLGTKFLLRTRENVNDYEITNQADIANSTFN
ASRPTKVLIHGFVLGGKEIAPWVPTMADAILGKENVNVILVNWVQGAKTQYDQAVANT
RMVGAQIHRLINRLSRKHGAKANDFHLIGFSLGSHVAGFAGKASIKSGKQLGRISGLDPA
NPAFNSNSSEVRLDRTDAKFVDVIHTDIRTILHISSGMNRSLGHIDFWPNGGESQPGCHN
WNGGIFQGFQDMAVCDHLRAPQLYIASIITTLPMIGYRCPNYDSFRRGTCLNCRGKVGSH
KNNRCAVMGYYAEPPRGSRKDAQVDYYLDTSDTSPYANRHYQVLIHWRKVPGKTTEE
GQRASLFLRLHGEYMSSEPRKLFLYKDLSTKYYHVYHNRTARFMITLPWDQDLGDLKK
VTFWWQRQTCWHYIGCKNDENIFVKRIRVLDAVEQKRYTFITPERLEHERVAPREILPND
KVEFVARQHHPKHKNSRSGRKGVEEADIVEPLLRQPMIVRRDTSSSSTSQSLSN

**Figure B: Open Reading Frame translation of PL002**

**Table C: Specific activity of P-Nitrophenol butyrate-based lipase reactions**

| Enzyme | Taxonomy | Temperature (°C) | Activity (U/mg) | Protein extraction | Reference |
|---|---|---|---|---|---|
| ***Malbranchea cinnamomea*** | Fungus | 40 | 263.3 ± 4.6 | Recombinant | (Duan et al. 2019) |
| ***Alkalispirillum sp. NM-ROO2*** | Bacteria | 52 °C | 38.1 ± 1.7 | Recombinant | (Mesbah 2019) |
| ***Aspergillus niger*** | Fungus | 40 | 0.22 | Recombinant | (V. C. Badgujar et al. 2017) |
| ***Aspergillus niger*** | Fungus | 30 | 1293 | Recombinant | (Cong et al. 2019) |
| ***Aspergillus oryzae*** | Fungus | 45 | 7.9 | Extracted | (Q. Li et al. 2021) |
| ***Bacillus licheniformis*** | Bacteria | 90 | 15916 | Extracted | (Ugras 2017) |
| ***Bacillus licheniformis – cold tolerant version*** | Bacteria | 35 | 93.37 ± 2.45 | Extracted | (Zhao et al. 2021) |
| ***Bacillus lipase*** | Bacteria | 30 | 79.42 | Extracted | (Jain and Mishra 2015) |
| ***Bacillus subtilis* lipase A** | Bacteria | 25 | 9.4 ± 0.5 | Recombinant | (Zhou et al. 2019) |
| ***Burkholderia cepacia*** | Bacteria | 25 | 29.11 | Recombinant | (K. C. Badgujar and Bhanage 2015) |
| ***Callosobruchus maculatus (beetle)*** | Animalia | 37 | 0.36 | Extracted | (Malaikozhundan and Vinodhini 2018) |
| ***Candida antarctica*** | Fungus | 60 | 22.5 ± 0.5 | Recombinant | (Carniel et al. 2017) |
| ***Candida cylindracea*** | Fungus | 40 | 0.83 | Recombinant | (V. C. Badgujar et al. 2017) |
| ***Candida rugosa*** | Fungus | 40 | 0.28 | Recombinant | (V. C. Badgujar et al. 2017) |
| ***Candida rugosa*** | Fungus | 25 | 12.15 ± 0.12 | Recombinant | (de Morais et al. 2016) |

| | | | | | |
|---|---|---|---|---|---|
| ***Ectomyelois ceratoniae (locust)*** | Animalia | 30 | 0.4 | Extracted | (Ranjbar et al. 2015) |
| ***Geotrichum candidum (human microbiome)*** | Fungus | 25 | 11.48 ± 0.13 | Recombinant | (de Morais et al. 2016) |
| ***Lactobacillus rhamnosus*** | Bacteria | 37 | 0.81 | Extracted | (Manasian et al. 2020) |
| ***Marinactinospora thermotolerans*** | Bacteria | 37 | 139.24 | Recombinant | (Deng et al. 2016) |
| ***Mucor javanicus*** | Fungus | 40 | 0.21 | Recombinant | (V. C. Badgujar et al. 2017) |
| ***Mycobacterium tuberculosis*** | Fungus | 37 | 35.71 | Recombinant | (Lin et al. 2017) |
| ***Oryza sativa (rice)*** | Plantae | 35 | 4.73 | Extracted | (Chen et al. 2019) |
| ***Paenibacillus pasadenensis*** | Bacteria | 50 | 1165.57 | Recombinant | (Gao et al. 2018) |
| ***Pancreatic like lipase, Salpa thompsoni (Zooplankton)*** | Animalia | 20 | 3.16 ± 0.14 | Recombinant | This study |
| ***Penicillium expansum*** | Fungus | 35 | 26.5 | Recombinant (yeast) | (L. Tang et al. 2015) |
| ***Proteus sp. NH 2-2 – Ecoli expressed*** | Bacteria | 20 | 24.16 ± 1.13 | Recombinant | (Shao et al. 2019) |
| ***Rhizomucor miehei*** | Fungus | 50 | 0.177 | Recombinant | (Yildirim et al. 2019) |
| ***Rhizomucor miehei*** | Fungus | 40 | 0.8 | Recombinant | (V. C. Badgujar et al. 2017) |
| ***Rhizopus oryzae*** | Fungus | 40 | 112.7 | Recombinant | (Zhao et al. 2019) |
| ***Rhodothermus marinus*** | Bacteria | 60 | 2073.3 | Recombinant | (Memarpoor-Yazdi, Karbalaei-Heidari, and Doroodmand 2018) |

| | | | | | |
|---|---|---|---|---|---|
| ***Ricinus communis L – (Castor oil plant)*** | Plantae | 40 | 47.902 ± 2.30 | yeast expressed | (Y. Li et al. 2021) |
| ***Stomolophus meleagris (Jellyfish)*** | Animalia | 55 | 0.80 ± 0.30 | Extraction | (Martínez-Pérez et al. 2020) |
| ***Thermomyces lanuginosus*** | Fungus | 30 | 2.74 | Extracted | (K. Tian et al. 2017) |
| ***Thermomyces lanuginosus*** | Fungus | 30 | 6 | Recombinant | (Noro et al. 2020) |
| ***Thermotoga maritima*** | Bacteria | 70 | 64.3 | Recombinant | (R. Tian et al. 2015) |
| ***Vicia faba (faba bean)*** | Plantae | 37 | 4500 | Extracted | (Lampi et al. 2020) |
| ***Yarrowia lipolytica lipase 2*** | Fungus | 40 | 140 | Recombinant | (Guilong Wang et al. 2015) |
| ***Yersinia enterocolitica*** | Bacteria | 37 | 1093.4 | Extracted | (Ji et al. 2015) |
